# β-catenin interacts with the TAZ1 and TAZ2 domains of CBP/p300 to activate gene transcription

**DOI:** 10.1101/2022.08.30.505852

**Authors:** Alexandra D. Brown, Connor Cranstone, Denis J. Dupré, David N. Langelaan

**Affiliations:** Department of Biochemistry & Molecular Biology, Dalhousie University, Halifax, NS, B3H 4R2, Canada; Department of Pharmacology, Dalhousie University, Halifax, NS, B3H 4R2, Canada

**Author notes:** To whom correspondence should be addressed: David N. Langelaan, Department of Biochemistry & Molecular Biology, Dalhousie University, Halifax, NS, B3H 4R2, Canada.

**Keywords:** β-catenin, transactivation domain, CBP/p300

## Abstract

The transcriptional co-regulator β-catenin is a critical effector of the canonical Wnt-signalling pathway, which plays a crucial role in regulating cell fate and maintaining tissue homeostasis. Deregulation of the Wnt/β-catenin pathway is characteristic in the development of major types of cancer, where accumulation of β-catenin promotes cancer cell proliferation and renewal. β-catenin gene expression is facilitated through recruitment of co-activators such as histone acetyltransferases CBP/p300; however, the mechanism of their interaction is not fully understood. Here we investigate the interaction between the C-terminal transactivation domain of β-catenin and CBP/p300. Using a combination of pulldown assays, isothermal titration calorimetry, and nuclear resonance spectroscopy we determine the disordered C-terminal region of β-catenin binds promiscuously to the TAZ1 and TAZ2 domains of CBP/p300. We then map the interaction site of the C-terminal β-catenin transactivation domain onto TAZ1 and TAZ2 using chemical-shift perturbation studies. Luciferase-based gene reporter assays indicate Asp750-Leu781 is critical to β-catenin gene activation, and mutagenesis revealed that acidic and hydrophobic residues within this region are necessary to maintain TAZ1 binding. These results provide a mechanistic understanding of Wnt/β-catenin gene regulation that underlies cell development and provide a framework to develop methods to block β-catenin dependent signalling in the future.

## 1. Introduction

Wnt signalling is a highly conserved signal transduction pathway that is critical to embryotic development and cell regulation throughout a person’s lifetime (1–3). Binding of the Wnt ligand to its receptor triggers two types of molecular cascades, canonical and non-canonical, which are differentiated based on the involvement of β-catenin (4). In the canonical Wnt pathway, β-catenin is responsible for mediating the transcriptional response (5). Upon activation, stabilized β-catenin enters the nucleus where it binds and modulates the actions of DNA-bound transcription factors such as t-cell factor/lymphoid enhancer factor (TCF/LEF), which require β-catenin for Wnt-targeted gene expression (5). These Wnt/β-catenin-target genes are essential in controlling numerous cellular processes including proliferation, differentiation, migration, and cell survival (6,7). Consequently, mutations to the Wnt/β-catenin signalling pathway are characteristic in many major types of cancer, where enhanced accumulation of nuclear β-catenin leads to tumor formation and cancer progression (8, 9).

β-catenin is composed of a central armadillo repeat domain (ARM) spanning residues Asn138-Asp665 that is flanked by N- and C-terminal sequences Met1-Val137 (βCAT_NTERM_) and Lys666-Asp781 (βCAT_CTERM_), respectively (Fig. 1) (10). Together these domains allow β-catenin to act as a scaffold for multi-protein complexes involved in transcription activation. The β-catenin ARM domain consists of 12 armadillo repeats that form a super-helical positively charged groove and provides the main interaction site for a multitude of transcription factors (11). βCAT_NTERM_ and βCAT_CTERM_ facilitate protein-protein interactions with transcriptional co-regulators, and are critical to β-catenin function as they both contain potent transactivation domains (TAD1/2) (12–14). This in part is attributed to the structurally disordered nature of these domains which allows β-catenin the flexibility to recruit co-activators to regulatory complexes associated with DNA (15). These include chromatin remodelling ATPases ISWI and BRG-1, and the homologous histone acetyltransferases CREB-binding protein (CBP) and E1A-binding protein (p300), which directly interact with β-catenin at gene promoters to enhance transcription (16–18). Interactions of CBP/p300 with β-catenin synergistically activate β-catenin/Wnt transcription, and lowering CBP/p300 levels inhibits growth and formation of colon carcinomas caused by β-catenin mutation (19).

**Fig. 1.**
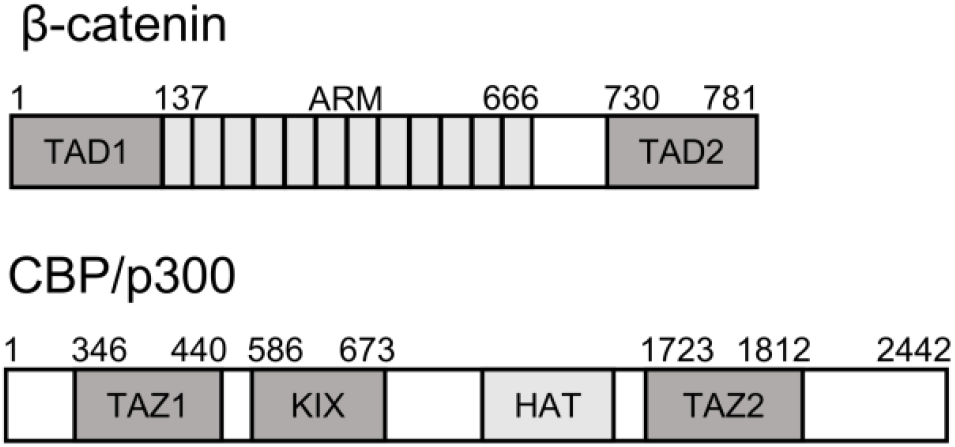
Domain architecture of β-catenin and CBP/p300. Schematic illustrating the domains of β-catenin including the central armadillo repeat domain (ARM), N-terminal, and C-terminal transactivation domains (TAD1/2). Domains of CBP/p300 include the histone acetyltransferase domain (HAT), and protein interacting domains KIX, TAZ1, and TAZ2. Numbered residues indicate domain boundaries in accordance with native sequence of p300.

CBP/p300 are critical histone acetyltransferases that accumulate at thousands of different human gene promoters and provide interaction sites for hundreds of known transcription regulators (20–22). Recruitment of CBP/p300 enhances target gene expression both by modulating chromatin structure through histone acetylation and by acting as an intermediary between transcription factors and transcriptional machinery (23). CBP/p300 are large proteins with seven globular domains, including the histone acetyltransferase (HAT) domain, which modifies histones and loosens chromatin thereby improving the ability of the associated DNA to be transcribed (24). CBP/p300 also has several protein interaction domains that are separated by long intrinsically disordered regions (Fig. 1). This includes the kinase inducible domain (KIX), which is recruited by the activation domains of transcription factors including MLL, BRCA1 and c-Myb (25–27). Additionally, the transcription adaptor zinc finger domains TAZ1 and TAZ2 facilitate similar interactions with the transactivation domains of HIF-1α, FOXO3, STAT1, and STAT2 (28–30).

Despite the importance of Wnt signalling in both health and disease and considerable evidence that interactions with CBP/p300 mediate Wnt signalling, the molecular mechanisms of how β-catenin interacts with CBP/p300 are not fully understood (16, 31, 32). Here, we characterize a direct interaction between TAD2 of β-catenin with the TAZ1 and TAZ2 domains of CBP/p300. We identify the residues involved in these interactions using nuclear magnetic resonance spectroscopy (NMR) and quantify their affinities using isothermal titration calorimetry (ITC). Finally, we used mutagenesis studies to evaluate the importance of certain motifs in modulating TAZ1-mediated CBP/p300 recruitment.

## 2. Materials and methods

### 2.1 Plasmid preparation

A pET28a plasmid containing full-length human β-catenin was gifted from Randall Moon (Addgene plasmid # 17198) and various fragments of the coding region (residues 1-137 (NTERM), 666-750, 666-781 (CTERM), 730-781 (TAD2), and 750-781) were amplified by PCR and cloned into a pET21b vector using *Eco*RI, *Bam*HI and *Xho*I restriction enzymes, downstream of sequences encoding for a hexahistidine tag, the B1 domain of *Streptococcus* protein G (GB1), and a tobacco etch virus (TEV) protease cleavage site to create pGB1-βCAT plasmids. Site-directed mutagenesis of pGB1-βCAT_CTERM_ was performed using sequence and ligation independent cloning (33), where various substitutions (D751A, D761A, D764A, F777A) and deletions (Δ749-753, Δ757-761, Δ761-765, Δ776-779) were incorporated into primers. Plasmids used for mammalian one-hybrid experiments included segments of the β-catenin coding region cloned into pCMV-GAL4 vector (gifted by Liqun Luo (Addgene plasmid # 24345)) to create pGAL4-βCAT plasmids. Plasmids coding for TAZ1 (residues 346–440 of CBP), TAZ2 (residues 1723-1812 of p300 with four stabilizing mutations C1738A, C1746A, C1789A, C1790A), and KIX (residues 586–673 of CBP) are the same as those previously used (34). The validity of all constructs was verified by DNA sequencing.

### 2.2 Protein expression and purification

pGB1-βCAT plasmids were transformed into chemically competent *E. coli* BL21 (DE3) and ampicillin (100 µg/mL) was added to all media for selection. Cultures were grown in LB media, ^15^N, or ^15^N/^13^C enriched M9 Minimal Media (35) at 37 °C to an optical density at 600 nm of 0.6-0.8, following which recombinant protein expression was induced with the addition of isopropyl β-D-1-thiogalactopyranoside (IPTG; 0.5 mM). Cultures were incubated for 4 hr at 37 °C and then harvested by centrifugation. Cell pellets for GB1-βCAT constructs were resuspended in 30 mL denaturing lysis buffer (20 mM Tris-HCl pH 8.0, 250 mM NaCl, 8 M urea), lysed by sonication, clarified by centrifugation, and purified by Ni^2+^ affinity chromatography (IMAC Sepharose, Cytiva), with protein refolding occurring on the column with the addition of lysis buffer lacking urea. The eluent was then dialyzed overnight against native buffer (20 mM Tris-HCl pH 8.0, 50 mM NaCl, 5 mM βME) at 4 °C. If required, TEV protease (150 µg) was added to remove the affinity tag and cleaved βCAT peptides were then separated from GB1 by Ni^2+^ affinity and ion exchange chromatography (Q Sepharose, Cytiva).

Purification of GB1-TAZ2 was carried out in same manner as GB1-βCAT proteins, with addition of 100 μM ZnCl_2_ added to all buffers. Following Ni^2+^ affinity chromatography, the eluent was reduced with the addition of β-mercaptoethanol (βME, 50 mM), diluted two-fold with native buffer containing 1 mM ZnCl_2_, and then incubated with 200 U thrombin overnight at 4 °C. Cleaved TAZ2 samples were diluted three-fold with a low-salt buffer (20 mM Tris-HCl pH 8, 5 mM βME, 10 μM ZnCl_2_), and purified by ion exchange chromatography (SP Sepharose, Cytiva). Purification of TAZ1 and KIX occurred as previously described (34), with additional semi-preparative reverse phase high-performance liquid chromatography purification for TAZ1 carried out on a C_8_ column using a water: acetonitrile gradient (Zorbax, Agilent Technolgies). All protein purification fractions were monitored by SDS-PAGE.

### 2.3 Pulldown assay

GB1 and GB1-βCAT fusion proteins (20 nmoles) were immobilized onto 20 μL IgG agarose beads (Cytiva) in pulldown buffer (20 mM Tris-HCl pH 8, 25 mM NaCl, 5 mM βME, and 10 μM ZnCl_2_). After washing away excess protein the beads were incubated with 20 nmoles of KIX, TAZ1, or TAZ2 for 30 min. Following incubation, beads were washed three times with pulldown buffer, resuspended in Laemmli buffer, and analyzed by SDS-PAGE. All experiments were performed in duplicate, statistical significance was measured using one-way ANOVA and Dunnett’s multiple comparison test to GB1 control, with a significance threshold p-value ≤ 0.01 and variation reported as standard error.

### 2.4 Isothermal titration calorimetry

Experiments were performed at 30 °C in 20 mM Tris-HCl pH 8.0, 25 mM NaCl, 5 mM βME, and 1 μM ZnCl_2_ using a VP-ITC microcalorimeter (MicroCal). TAZ1 or TAZ2 (100-200 μM) was injected into a calorimetric reaction cell containing 8-20 μM βCAT peptides. ITC experiments were collected in duplicate with 30 increments of 10 μL injections at 300 sec equilibration intervals. Thermograms were fit to the appropriate one-site or two-site binding models using MicroCal Origin 7.5 software.

### 2.5 NMR spectroscopy

Unless otherwise noted. all NMR spectra of were collected in 20 mM MES pH 6.2, 100 mM NaCl, 5 mM DTT, 5% D_2_O, at 25 °C on a Bruker Avance III 700 MHz spectrometer equipped with a cryogenically cooled probe at the National Research Council Institute for Marine Biosciences (NRC-IMB, Halifax, NS). Resonance assignments of ^13^C/^15^N-labelled βCAT_CTERM_ (500 µM) were determined by interpreting ^1^H-^15^N HSQC, HNCACB, CBCACONH, H(CCO)NH, (H)C(CO)NH, HNCO, and HN(CA)CO experiments. Chemical shift assignments of backbone resonances for TAZ1 and TAZ2 were determined by analyzing triple resonance spectra that were collected at 600 MHz (Varian INOVA, Queen’s University, Kingston ON) from samples of ^13^C/^15^N-labelled protein (TAZ1: 650 µM TAZ1, 20 mM MES pH 6.5, 5 mM βME, 5% D_2_O; TAZ2: 358 µM TAZ2, 20 mM MES pH 6.5, 5 mM βME, 100 µM ZnCl_2_, 5% D_2_O) at 25 °C and 15 °C, respectively. NMR data were processed using NMRPipe (36) and analyzed using CcpNmr Analysis (37).

To assess binding between β-catenin and domains of CBP/p300, ^1^H-^15^N HSQC spectra were collected of 100 µM ^15^N-labeled βCAT_CTERM_ or βCAT_750-781_ in the absence and presence of up to 400 μM unlabelled TAZ1 or TAZ2. Chemical shift changes (Δδ) upon addition of TAZ1 or TAZ2, were quantified for each residue as Δδ = [(0.17ΔδN)^2^ + (ΔδHN)^2^]^1/2^ (38). For mapping experiments, ^1^H-^15^N HSQC spectra were collected of 100 µM ^15^N-labeled TAZ1 or TAZ2 titrated with up to 400 μM unlabelled βCAT_TAD2_ or βCAT_750-781_ in NMR buffer with 25 mM NaCl at 35 °C. Residues that experienced Δδ greater than the mean Δδ or mean Δδ + 1 standard deviation, were mapped onto the surface of the 3D crystal structure of TAZ1 and TAZ2 (PDBID: 1U2N and 1F81, respectively) (39, 40). PyMOL (Schrödinger Inc.) was used to interpret structures and generate figures. Chemical shift assignments for βCAT_CTERM_ are deposited into the BioMagResBank as accession number 51601 (41).

### 2.6 Luciferase-based transactivation assays

HEK 293A cells were cultured in Dulbecco’s modified Eagle’s medium supplemented with 10% fetal bovine serum at 37°C with 5% CO_2_. For reporter assays, cells were seeded into 24-well plates and transfected the following day at 70-80% confluency using jetPRIME transfection reagent. A total of 0.5 µg of plasmid was transfected per well including 0.35 µg luciferase reporter (p5xGAL4-luc), 0.05 µg internal control (pCMV-Renilla) and 0.1 µg of pGAL4-βCAT expression plasmid. Luciferase activity was determined 18-24 hr post-transfection from cell lysate using the Dual-Luciferase Reporter Assay System (Promega) according to the manufacturers protocols. All luciferase values are an average of *Renilla-*normalized luminescence and represent at least three independent transfections. Values are expressed as fold activation, the ratio of renilla-normalized luciferase activity over pCMV-GAL4 negative control, where error bars indicate standard error. Statistical significance was measured using one-way ANOVA and Dunnett’s multiple comparison test relative to GAL4 control, with a significance threshold p-value ≤ 0.01.

## 3. Results

### 3.1 The C-terminal transactivation domain of β-catenin is intrinsically disordered

NMR spectroscopy was first used to investigate the properties of βCAT_CTERM_ (residues 666-781). The ^1^H-^15^N HSQC spectrum of ^13^C/^15^N-labelled βCAT_CTERM_ was high quality and had the expected number of resonance peaks with sharp linewidths and low chemical shift dispersion. This is consistent with βCAT_CTERM_ being intrinsically disordered and with the absence of this region from the crystal structure of full-length β-catenin (10). Standard triple resonance experiments were utilized to manually assign to assign 115/118 residues and 94% of the backbone resonances of βCAT_CTERM_ (Fig. 2A). Secondary structure propensity analysis (42), indicates that there are no extended regions of sequence with a secondary structure propensity score magnitude > 0.3, suggesting that βCAT_CTERM_ does not have significant residual α-helix or β-sheet secondary structure (Fig. 2B).

**Fig. 2.**
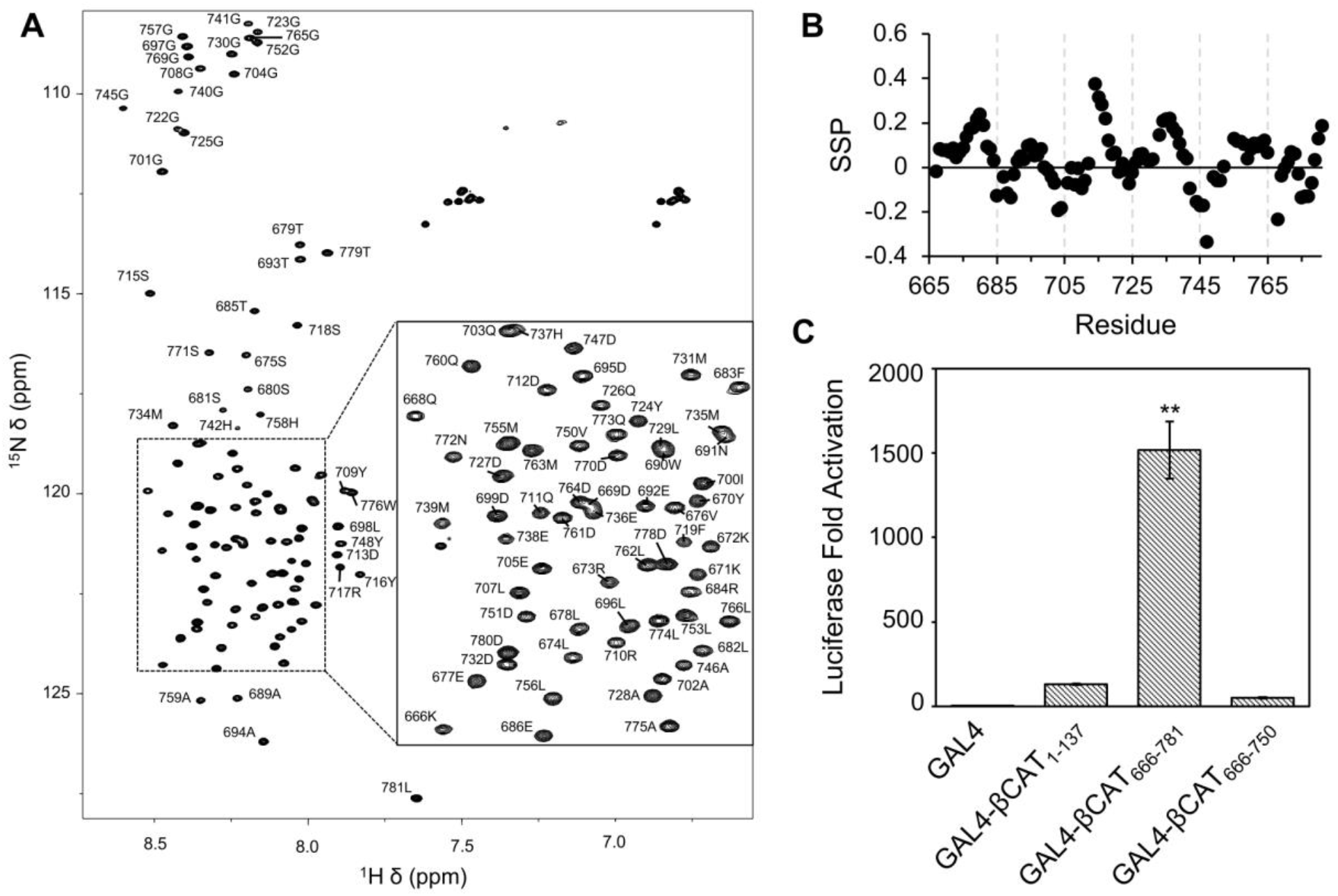
The C-terminal transactivation domain of β-catenin is disordered (A) ^1^H-^15^N HSQC of βCAT_CTERM_ with residue assignments of peaks indicated. (B) Secondary structure propensity (SSP) values per residue of βCAT_CTERM_ calculated from Cα and Cβ chemical shifts. (C) Luciferase-based mammalian one-hybrid transactivation assay performed in HEK 293A cells using the indicated pGAL4-βCAT fusion proteins. Statistical significance set at p-value ≤ 0.01** by one-way ANOVA and Dunnett’s multiple comparison test compared to pGAL4 control, variation is reported as standard error.

To better define the boundaries of the C-terminal β-catenin transactivation domain, the ability of various regions of β-catenin to influence transcription was tested using a luciferase-based one-hybrid gene reporter assay. HEK 293A cells were co-transfected with a luciferase reporter and a of mammalian expression plasmids encoding for a region of β-catenin (residues 1-137 (NTERM), CTERM, and 666-750) fused to a GAL4 DNA-binding domain. While some activity was observed for the N-terminal region of β-catenin, the C-terminal region of β-catenin demonstrated potent transcription activation of the luciferin reporter with 1500-fold more activation than GAL4 alone (Fig. 2C). Transfection with pGAL4-βCAT_666-750_ did not increase luciferase expression, indicating the functional importance of the C-terminal 31 amino acids for β-catenin transactivation.

### 3.2 The β-catenin transactivation domain 2 interacts with TAZ1 and TAZ2

The KIX, TAZ1, and TAZ2 domains of CBP/p300 are candidate interaction sites for intrinsically disordered activation domains of transcription regulators. To identify if any of these domains interact with the C-terminal transactivation domain of β-catenin, we completed a pulldown assay involving various segments of β-catenin (CTERM, 666-729, and 730-781 (TAD2)), expressed as fusion proteins with the B1 binding-domain of *Streptococcus* protein G (GB1). GB1-βCAT proteins were expressed in *E. coli*, purified, and then bound to IgG Sepharose beads and mixed with an equivalent amount of purified recombinant KIX, TAZ1, or TAZ2. SDS-PAGE analysis of bound protein indicates that βCAT_CTERM_ interacts with isolated TAZ1 and TAZ2 (Fig. 3). This association with TAZ1 and TAZ2 is maintained for βCAT_TAD2_ but lost for GB1-βCAT_666-729_. The KIX domain of CBP/p300 did not interact with any of the β-catenin constructs that were tested.

**Fig. 3.**
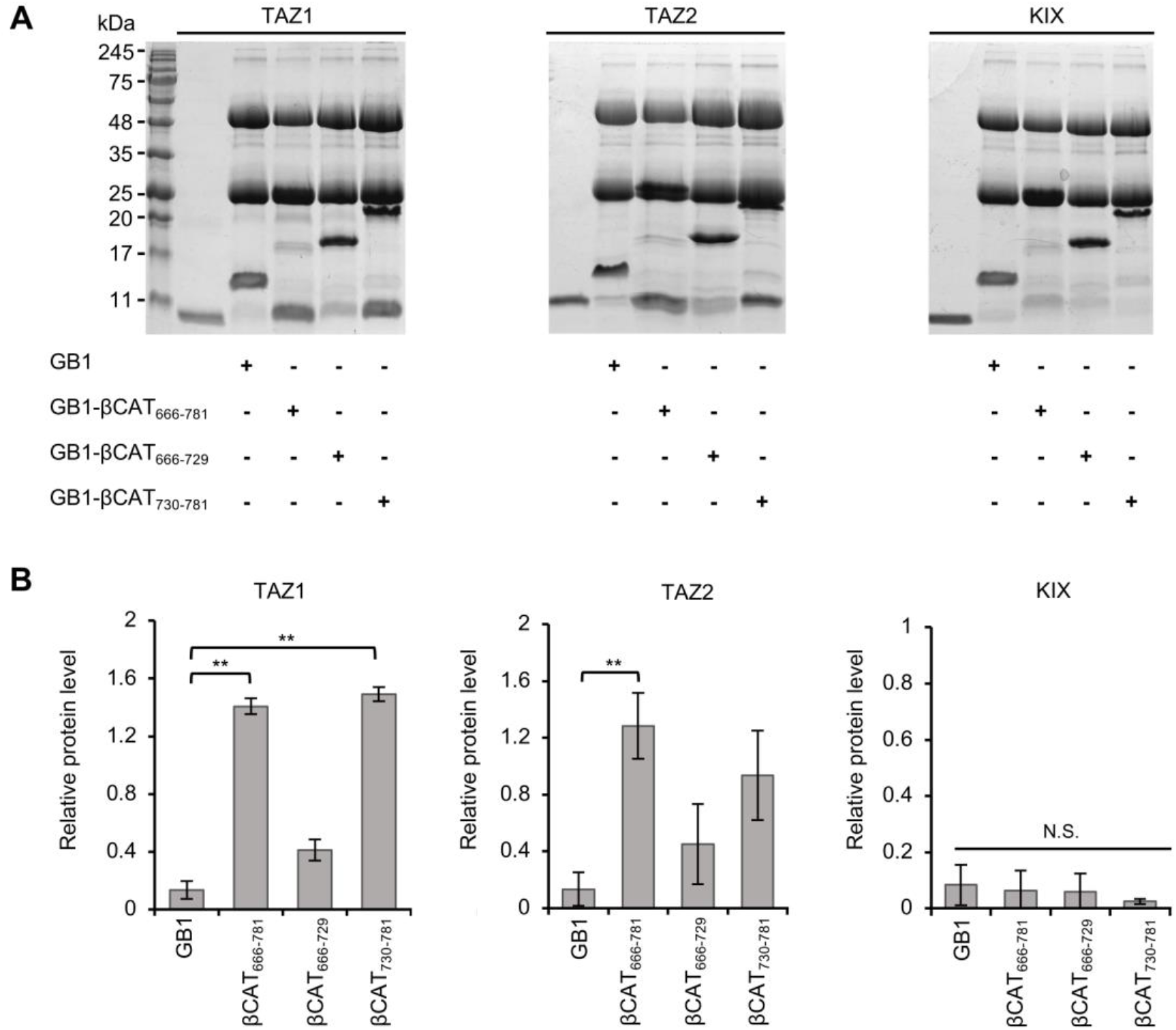
βCAT_CTERM_ binds the TAZ1 and TAZ2 domains of CBP/p300. (A) 15% SDS-PAGE analysis of the ability of various GB1-βCAT constructs immobilized on IgG agarose beads to pulldown and interact with purified TAZ1, TAZ2, and KIX domains of CBP/p300. Left lanes show migration of each isolated CBP/p300 domain. (B) Quantitative analysis of protein pulldown presented in (A), statistical significance between protein pulldown levels of TAZ1, TAZ2, or KIX set at p-value ≤ 0.01** by one-way ANOVA and Dunnett’s multiple comparison test compared to GB1, error bars represent standard deviation.

### 3.3 Characterization of β-catenin:TAZ1 binding

To determine what residues of the β-catenin C-terminal are responsible for TAZ1 binding, a ^15^N-labelled βCAT_CTERM_ sample was prepared and titrated with unlabelled TAZ1 (Fig. 4A and full spectrum shown in Fig. S1). A subset of peaks shifted upon addition of TAZ1, indicating that there is a specific interaction between these proteins. Comparisons of chemical shift changes (Δδ, Fig. 4B) in the absence and presence of TAZ1 show residues Phe683-Ile700 and those C-terminal of Leu729 experience the largest chemical shift changes, with many residues between Asp750-Leu781 undergoing line broadening and not being visible upon addition of TAZ1. These changes indicate that TAZ1 interacts with these regions of β-catenin. ITC revealed that two different sites of βCAT_CTERM_ have affinity for TAZ1, since two-site binding was required to fit the collected thermograms when βCAT_CTERM_ was titrated into TAZ1 (Fig. 4C, K_d1_ = 14 ± 4.4 nM, K_d2_ = 0.65 ± 0.22 µM). In contrast, βCAT_TAD2_ shows one-site binding to TAZ1, with a dissociation constant of 0.38 ± 0.02 µM (Fig. 4D). Consistent with the pulldown data, a low-affinity one-site binding interaction is seen between βCAT_666-729_ and TAZ1 (Fig. S2A, K_d_ 67 ± 20 µM).

**Fig. 4.**
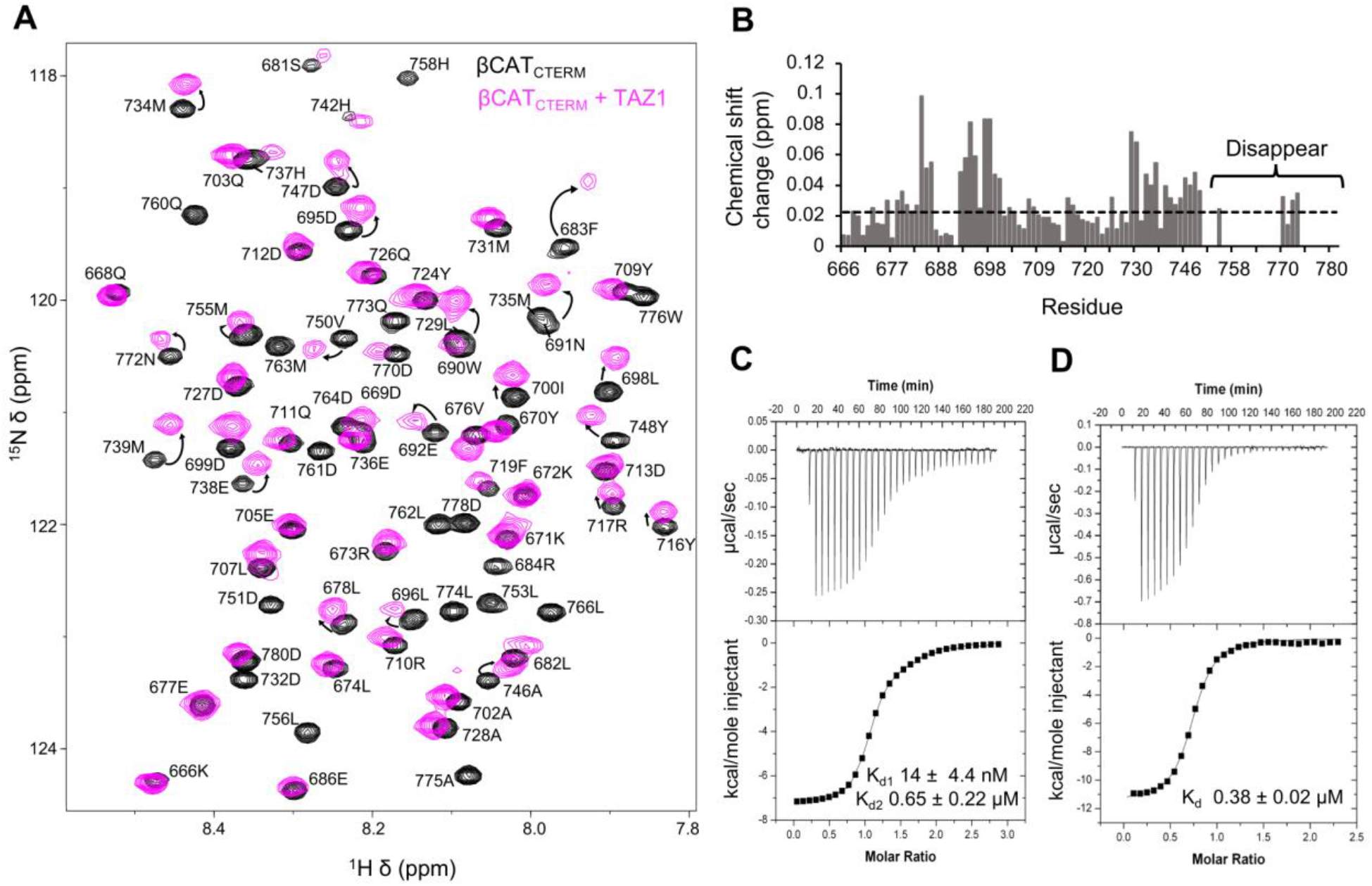
βCAT_CTERM_ binds TAZ1 with high-affinity. (A) ^1^H-^15^N HSQC of 100 µM ^15^N-labelled βCAT_CTERM_ overlayed in the absence (black) and presence of 400 µM purified recombinant TAZ1 (pink), annotated arrows indicate residues experiencing chemical shift upon addition of TAZ1. (B) Chemical shift changes (Δδ = [(0.17Δδ_N_)^2^+ (Δδ_HN_)^2^]^1/2^) that each residue of βCAT_CTERM_ experiences upon addition of TAZ1. Resonances that disappear due to line-broadening are noted. (C) Isothermal titration calorimetry thermogram of TAZ1 titrated into βCAT_CTERM_ fit to a two-site binding model yielding K_d1_ = 14 ± 4.4 nM and K_d2_ = 0.65 ± 0.22 µM. (D) ITC thermogram of TAZ1 titrated into βCAT_TAD2_ fit to a one-site binding model yielding K_d_ = 0.38 ± 0.02 µM.

The loss of signal for many residues between Asp750-Leu781 is likely due to chemical exchange induced by the interaction with TAZ1. βCAT_TAD2_ still binds TAZ1, albeit weaker than βCAT_TAD2_, with a dissociation constant of K_d_ 2.17 ± 0.07 µM. (Fig. S2B). To characterize this interaction, ^15^N-labelled βCAT_750-781_ was titrated with unlabelled TAZ1 and chemical shift changes were observed using NMR spectroscopy. With this smaller peptide all resonances remained visible when interacting with TAZ1 (Fig. S3), with Gly757-Asp761 and Trp776-Asp780 experiencing the largest chemical shift changes for any residue. This strong binding of motifs within βCAT_750-781_ to TAZ1 is consistent with deletion of this sequence ablating transactivation by GAL4-βCAT_CTERM_ (Fig. 2).

NMR-based chemical shift mapping studies were performed to define where on the TAZ1 surface βCAT_750-781_ binds (Fig. 5). Upon addition of βCAT_750-781_, a subset of resonances of ^15^N-labelled TAZ1 shifted, indicative of a specific interaction between these proteins (Figs. 5A, B). Mapping of the significantly perturbed residues onto the structure of TAZ1 (PDBID: 1U2N) (39), revealed βCAT_750-781_ binds to a shallow hydrophobic groove formed by the junction of the α1 and α4 helices of TAZ1 (Fig. 5C). In contrast, few significant chemical shift changes were observed on the opposing site of TAZ1 at helices α2 and α3.

**Fig. 5.**
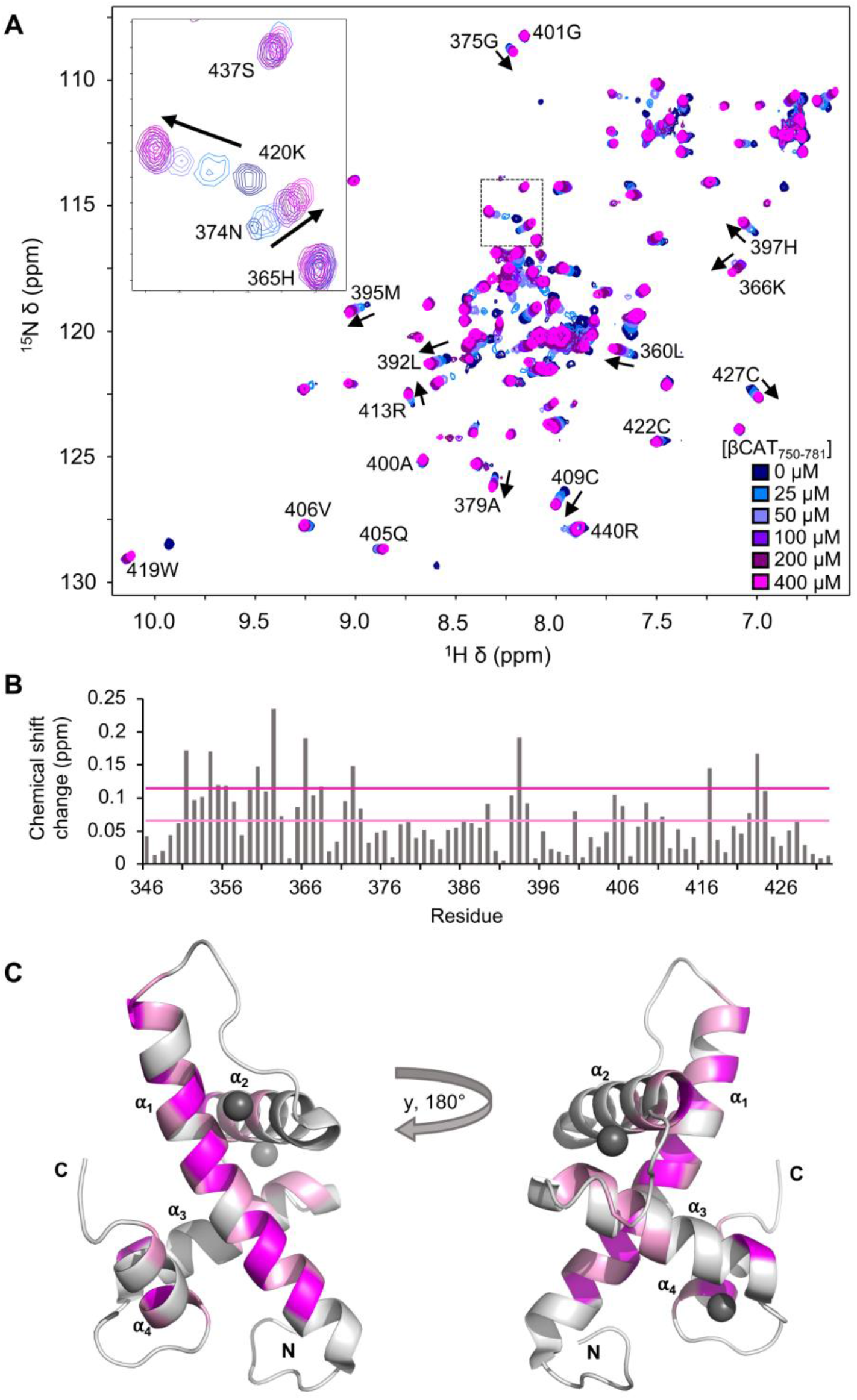
βCAT_750-781_ interaction site mapped onto TAZ1 surface. (A) Overlay of ^1^H-^15^N HSQC of 100 µM ^15^N-labelled TAZ1 titrated with 400 µM of unlabelled βCAT_750-781_. (B) Chemical shift changes (Δδ = [(0.17Δδ_N_)^2^+ (Δδ_HN_)^2^]^1/2^) that each residue of TAZ1 experienced upon addition of βCAT_750-781_ are calculated for each residue. (C) Residues with above the mean Δδ (pink) and Δδ + standard deviation (magenta) mapped onto the ribbon representation of TAZ1 (PDBID: 1U2N).

### 3.4 Characterization of β-catenin:TAZ2 binding

To characterize the βCAT_CTERM_:TAZ2 interaction, ^15^N-labelled βCAT_CTERM_ was titrated with unlabelled TAZ2 and a shift of a subset of β-catenin amide resonances was observed (Fig. 6A and full spectrum shown in Fig. S4). Quantification of NMR-chemical shift perturbations indicate that two regions from residues Pro714-Gly741 and Asp750-Leu781 underwent the most significant chemical shift changes (Fig. 6B). Like TAZ1, ITC analysis indicates that βCAT_CTERM_ binds TAZ2 at two different sites with lower affinities than TAZ1 (K_d1_ = 0.95 ± 0.04 µM, K_d2_ = 3.42 ± 0.2 µM; Fig. 6C), with βCAT_TAD2_ maintaining high affinity for TAZ2 (K_d1_ 0.65 ± 0.06 µM, K_d2_ 1.18 ± 0.43 µM; Fig. 6D).

**Fig. 6.**
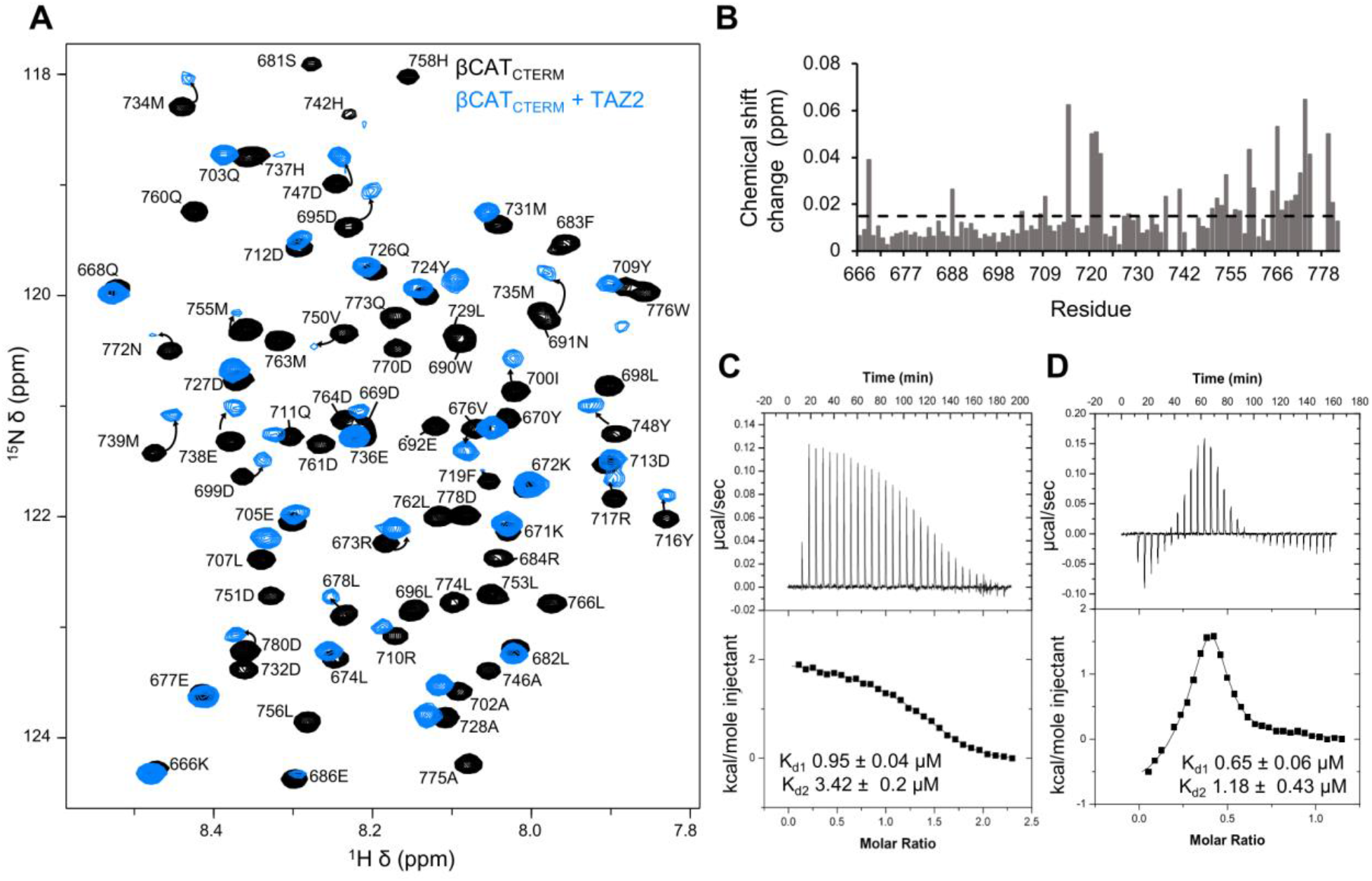
βCAT_CTERM_ binds TAZ2. (A) ^1^H-^15^N HSQC of 100 µM ^15^N-labelled βCAT_CTERM_ overlayed in the absence (black) and presence of 400 µM purified recombinant TAZ2 (blue), annotated arrows indicate residues experiencing chemical shift upon addition of TAZ2. (B) Chemical shift changes (Δδ = [(0.17Δδ_N_)^2^+ (Δδ_HN_)^2^]^1/2^) that each residue of βCAT_CTERM_ experienced upon addition of saturating amounts of TAZ2. (C) Isothermal titration calorimetry thermogram of TAZ2 titrated into βCAT_CTERM_ fit to a two-site binding model yielding K_d1_ = 0.95 ± 0.04 µM and K_d2_ = 3.42 ± 0.2 µM. (D) ITC thermogram of TAZ2 titrated into βCAT_TAD2_ fit to a two-site binding model yielding K_d1_ 0.= 65 ± 0.06 µM and K_d2_ = 1.18 ± 0.43 µM.

To determine the interaction site of βCAT_TAD2_ on the TAZ2 surface, ^15^N-labelled TAZ2 was titrated with unlabelled βCAT_TAD2_, and backbone amide chemical shift changes were monitored (Figs. 7A, B). Residues experiencing chemical shift changes larger than the average plus one standard deviation were mapped onto a previously determined structure of TAZ2 (PDBID: 1F81) (40). These findings indicate βCAT_TAD2_ binding is dispersed over the TAZ2 surface with select residues within α1-α4 experiencing large chemical shift changes, which is consistent with our finding that βCAT_TAD2_ interacts with TAZ2 at multiple sites (Fig. 7C).

**Fig. 7.**
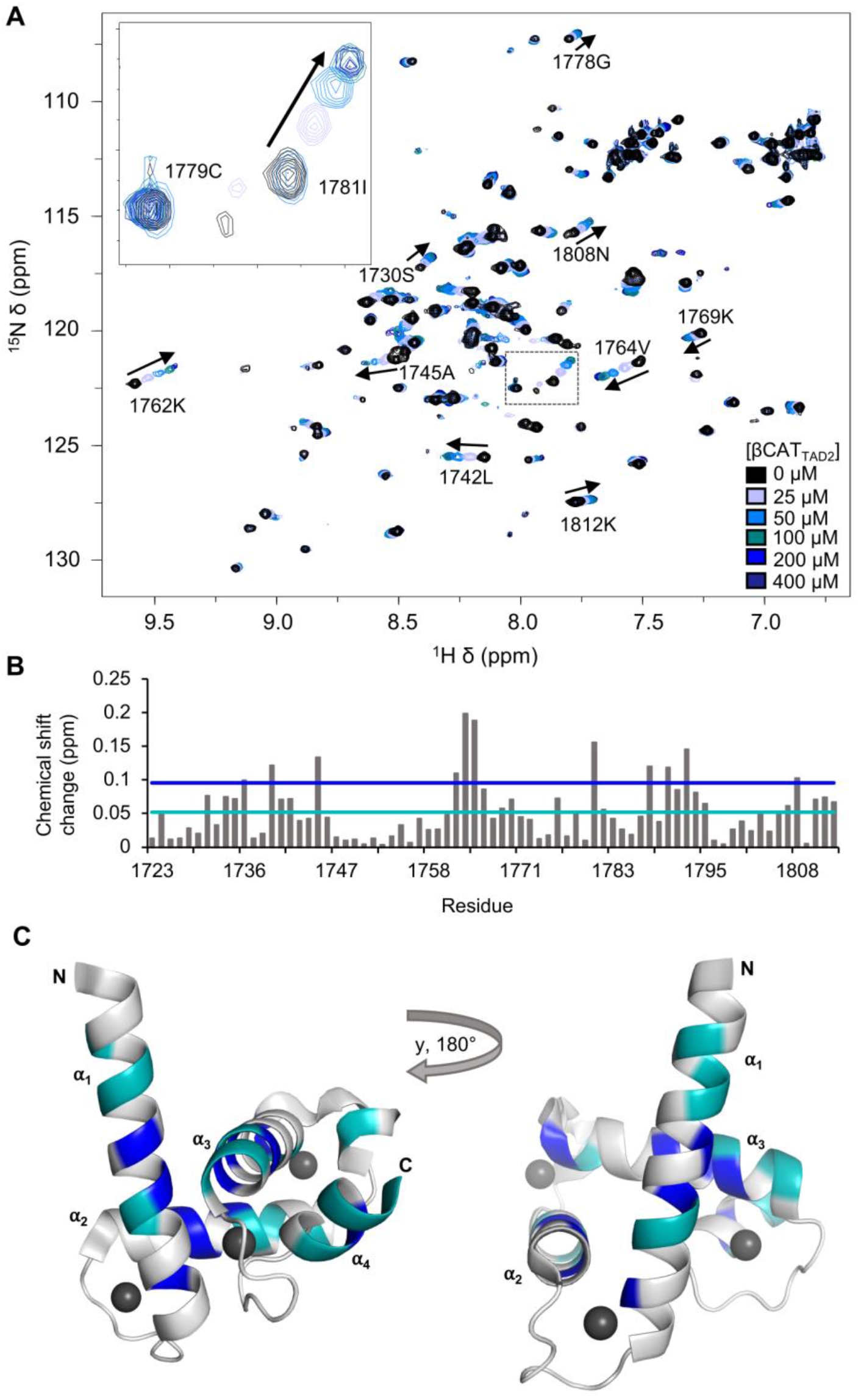
βCAT_TAD2_ interaction site mapped onto TAZ2 surface. (A) Overlay of ^1^H-^15^N HSQC spectra of 100 µM ^15^N-labelled TAZ2 titrated with 400 µM of unlabelled βCAT_TAD2_. (B) Chemical shift changes (Δδ = [(0.17Δδ_N_)^2^+ (Δδ_HN_)^2^]^1/2^) that each residue of TAZ2 experienced between free and bound to βCAT_TAD2_ calculated for each residue. (C) Residues with above the mean Δδ (blue) and Δδ + standard deviation (navy) mapped onto a ribbon representation of TAZ2 (PDBID: 1F81).

### 3.5 Hydrophobic and acidic residues mediate binding between β-catenin and TAZ1

Since β-catenin interacts more tightly and specifically with TAZ1 than TAZ2, we used mutagenesis to investigate which residues of β-catenin mediate this interaction. βCAT_750-781_ contains acidic and hydrophobic residues which are often important for binding CBP/p300, and many of these also experience above average shift perturbations upon addition of TAZ1 (Figs. 8A and S3). To test if these residues were critical to the β-catenin:TAZ1 interaction, site-directed mutagenesis was performed where acidic or hydrophobic residues of βCAT_CTERM_ that undergo large chemical shifts upon TAZ1 binding were substituted for alanine (D751A, D761A, D764A, F777A). ITC experiments determined these mutants bind TAZ1 with ∼20-40-fold lower affinity than wildtype βCAT_CTERM_, with measured dissociation constants of 0.39 ± 0.03 µM, 0.60 ± 0.06 µM, 0.32 ± 0.02 µM, and 0.55 ± 0.04 µM for D751A, D761A, D764A and F777A, respectively (Fig. 8B-E). Likewise, deletion of small motifs across βCAT_CTERM_ (Δ757-761, Δ761-765, Δ776-779) further decreased association with TAZ1 to the point that no binding was detectable by ITC (Fig. S5).

**Fig. 8.**
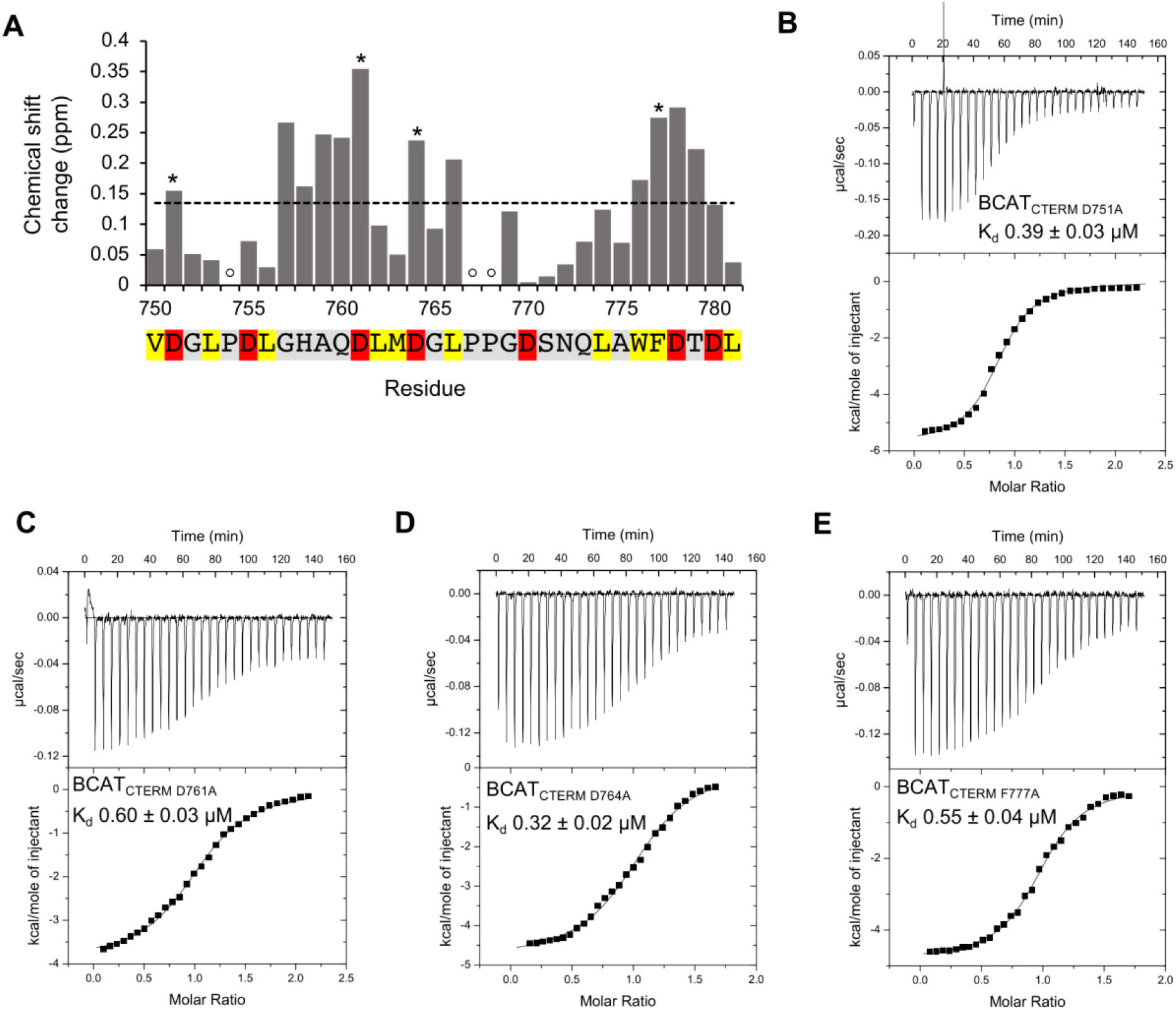
BCAT_CTERM_ binds TAZ1 though acidic and hydrophobic residues. (A) Chemical shift changes (Δδ = [(0.17Δδ_N_)^2^+ (Δδ_HN_)^2^]^1/2^) that each residue of βCAT_750-781_ experienced upon addition of saturating amounts of TAZ1. Residues with large Δδ that were selected for site-directed mutagenesis are denoted by *, while empty circles indicate proline residues that could not be monitored via ^1^H-^15^N HSQC. The sequence of βCAT_750-781_ is noted below the plot. (B-E) Isothermal titration calorimetry thermograms of TAZ1 titrated into β-CAT_CTERM_ mutants D751A, D761A, D764A, and F77A, fit to a one-site binding model yielding dissociation constants (K_d_) 0.55 ± 0.04 µM, 0.39 ± 0.03 µM, 0.60 ± 0.03 µM, and 0.32 ± 0.02 µM, respectively..

## 4. Discussion

β-catenin is a critical nuclear effector of the canonical Wnt-signalling pathway, however, aberrant activation causing hyperaccumulation of β-catenin is broadly implicated in many major types of cancer (43, 44). Nuclear β-catenin forms transcriptional complexes with co-activators such as CBP/p300, which together enhance the transcription of oncogenes driving cancer initiation and progression (45). Here we examine the recruitment of CBP/p300 by the C-terminal transactivation domain of β-catenin and elucidate fundamental molecular mechanisms of β-catenin-dependent gene transcription. We observe that βCAT_CTERM_ is intrinsically disordered and interacts with both the TAZ1 and TAZ2 domains of CBP/p300 with high affinity. This interaction is mediated by both acidic and hydrophobic amino acids within residues 750-781 of β-catenin, where mutagenesis of this region ablates transactivation potential and interactions with TAZ1.

Intrinsically disordered regions are abundant in transcription regulators, as their structural flexibility allows them to be highly dynamic and helps to facilitate promiscuous protein-protein interactions with a multitude of different binding partners. Promiscuity in binding of transcription regulators is key to the diversity of transcriptional complexes found at different gene promoters. The KIX, TAZ1, TAZ2 domains of CBP/p300 are candidate interaction sites for disordered activation domains of transcription regulators including CITED2, HIF1α, STAT2, and p53 (28, 30, 46, 47). While some activation domains associate with high specificity to particular CBP/p300 domains, others like the activation domain of tumor protein p53 have been shown to interact with promiscuity to TAZ1, TAZ2, and KIX with a broad range of affinity and specificity (48). Our observation that the β-catenin C-terminal transactivation domain interacts with both TAZ1 and TAZ2 in a redundant manner is consistent with this model of CBP/p300 recruitment. The binding promiscuity of the βCAT_CTERM_ is highlighted by how it contains two binding sites for both TAZ1 and TAZ2 (Figs. 4 and 6). This redundancy has also been observed in the E2A transcription factor, and may serve to increase the diversity of interactions that β-catenin can participate in at gene promoters (49).

TAZ1 has a tetrahedral shape and consists of four α-helices that create three hydrophobic grooves that bind activation domains (39). Using NMR-based chemical shift mapping studies, we determined that βCAT_750-781_ interacts with TAZ1 through an extended interface along helix α1, and in the shallow hydrophobic groove between helices α1 and α4 (Fig. 5), corresponding to the binding site of other activation domains including CITED2, HIF-1α, STAT2, and RelA (28, 30, 46, 50, 51). TAZ2 adopts a similar structure as TAZ1, but differentiated by the opposite orientation of helix α4 (29,30). Even though TAZ1 and TAZ2 have a similar topology, both domains differ in terms of sequence and binding specificity, with TAZ1 having deeper groves in its surface than TAZ2. Despite this, the flexibility of the β-catenin C-terminal transactivation domain can compensate for differences in topology and bind both TAZ1 and TAZ2 with high-affinity. NMR chemical shift mapping indicated that βCAT_TAD2_ binds TAZ2 through extended interactions across multiple helices (Fig 7), this is consistent with our ITC data that detected multiple binding sites between βCAT_TAD2_ and TAZ2, and similar to other transactivation domains like those of p53, which wrap around TAZ2 with multiple points of contact (52). Interestingly, p53 has been shown to compete with β-catenin for CBP/p300, which may be occurring through direct competition for TAZ1/2 domains at gene promoters (53).

Both TAZ1 and TAZ2 are basic proteins with a strongly electropositive surface that preferentially bind acidic motifs, where electrostatic interactions mediate activation domain recognition and specificity. Our findings demonstrate that deletion of acidic residues within the β-catenin transactivation domain significantly reduces βCAT_CTERM_:TAZ1 binding, highlighting the importance of electrostatic interactions for TAZ1 recognition (Fig. 8). TAZ1-binding favours longer extended disordered domains with multiple amphipathic motifs, which may explain why β-catenin point mutants that removed a single acidic or hydrophobic contact reduced but did not completely ablate the interaction between βCAT_CTERM_ and TAZ1. The association of β-catenin with CBP/p300 may also be regulated by post-translational modifications. CBP/p300-medaited acetylation of β-catenin is frequently disrupted by mutation in thyroid cancers, causing an increase in β-catenin activity (54). Additionally, numerous serine and threonine residues are present in the β-catenin transactivation domains. Similar to the p53 transactivation domains, which modulate interactions with CBP/p300 through phosphorylation (55), these residues may serve as potential phosphorylation sites that enhance the affinity of β-catenin for TAZ1 or TAZ2 by increasing the number of electrostatic interactions.

Overall, we have characterized how the C-terminal of β-catenin binds to both the TAZ1 and TAZ2 domains of CBP/p300, which provides mechanistic understanding of how β-catenin recruits CBP/p300 to mediate Wnt signalling. This work will aid future studies focused on interpreting the interplay between β-catenin and other transcription factors that bind TAZ1 and TAZ2, as well as developing molecules that disrupt β-catenin interactions with TAZ1 or TAZ2.

## Acknowledgements

We thank Dr. Stephen Bearne for providing access to an isothermal titration calorimeter and Mr. Ian Burton from the National Research Council Institute for Marine Biosciences (NRC-IMB) for assistance with NMR data acquisition.

