## Supplemental Material for "β-catenin interacts with the TAZ1 and TAZ2 domains of CBP/p300 to activate gene transcription"

**Contents:**

Supplementary figures S1-S5

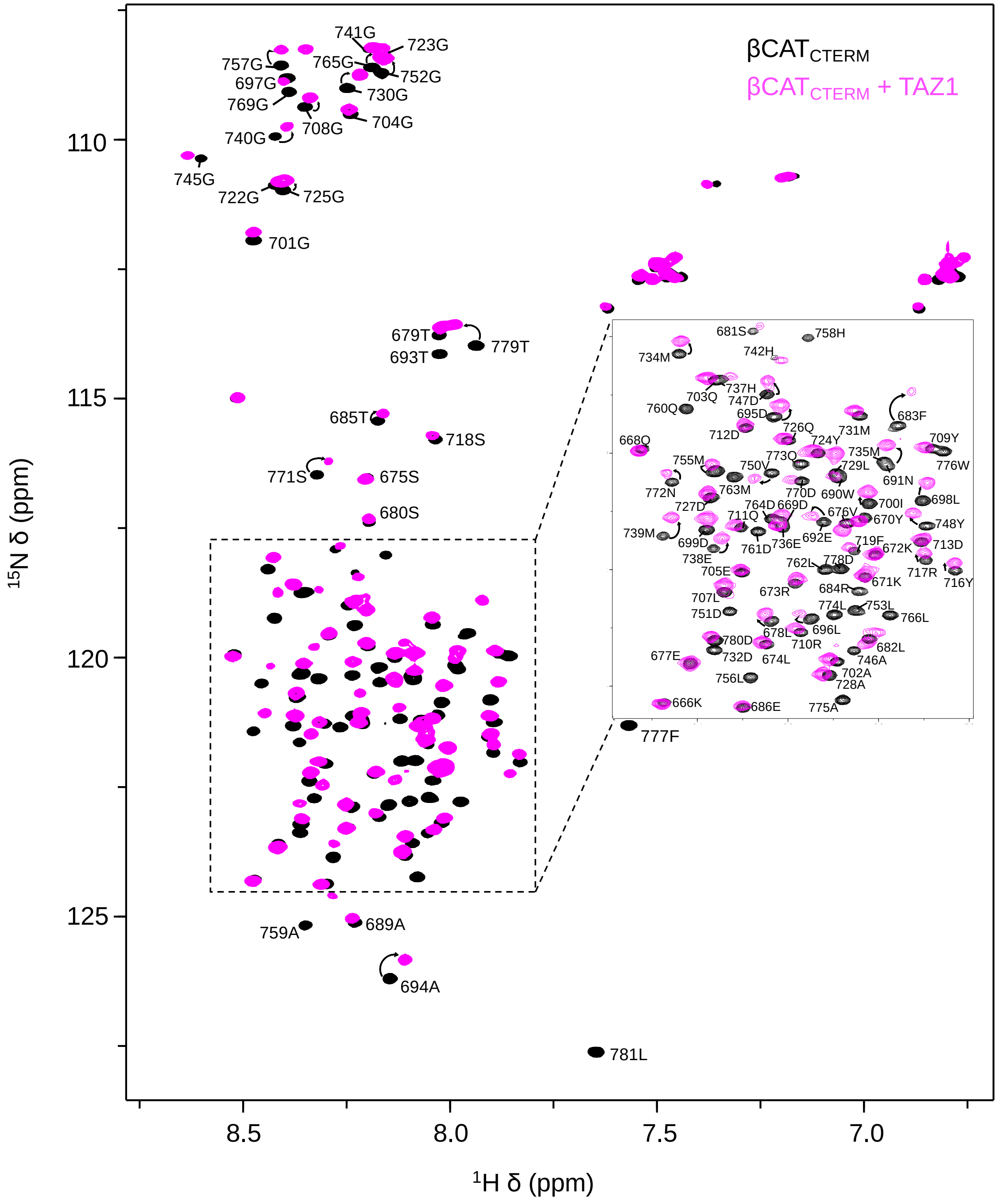


**Fig. S1.** $^{1}$H-$^{15}$N HSQC spectra of 100 μM $^{15}$N-labelled $\beta CAT_{CTERM}$ overlayed in the absence (black) and presence (pink) of 400 μM unlabeled TAZ1. Resonance assignments and direction of peak movement upon addition of TAZ1 are indicated.

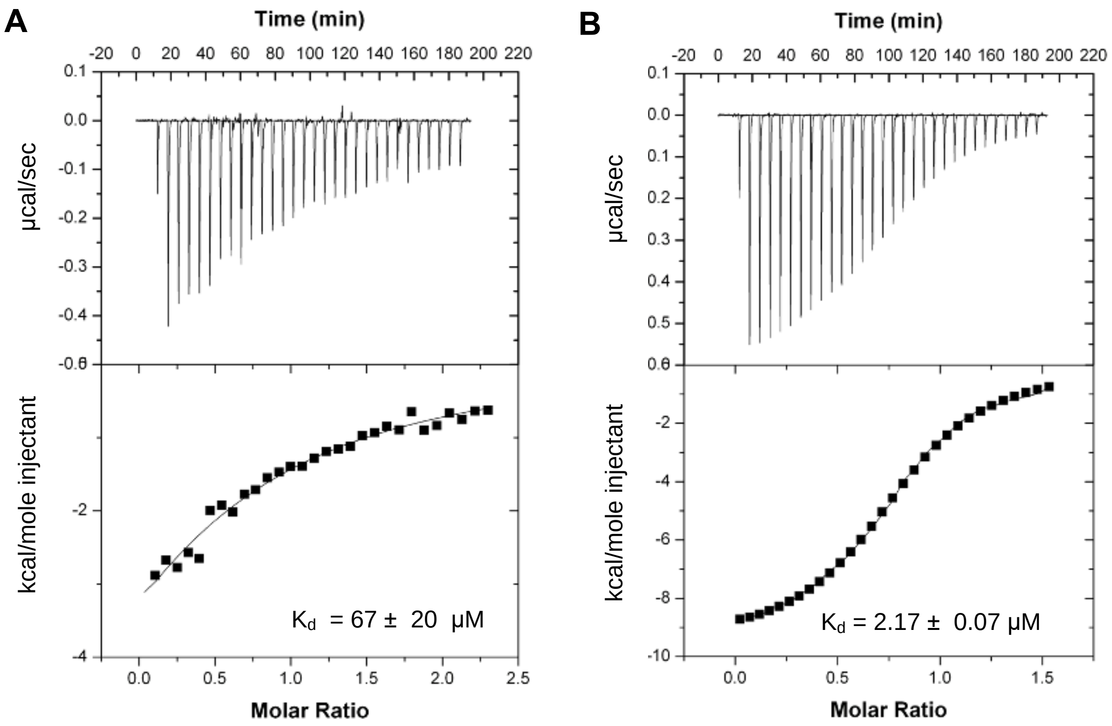


**Fig. S2.** (A) Isothermal titration calorimetry thermograms of 200 µM TAZ1 titration into 20 µM $\beta CAT_{666-729}$ ($K_d$ = 67 ± 20 µM) or (B) $\beta CAT_{750-781}$ ($K_d$ = 2.17 ± 0.07 µM)

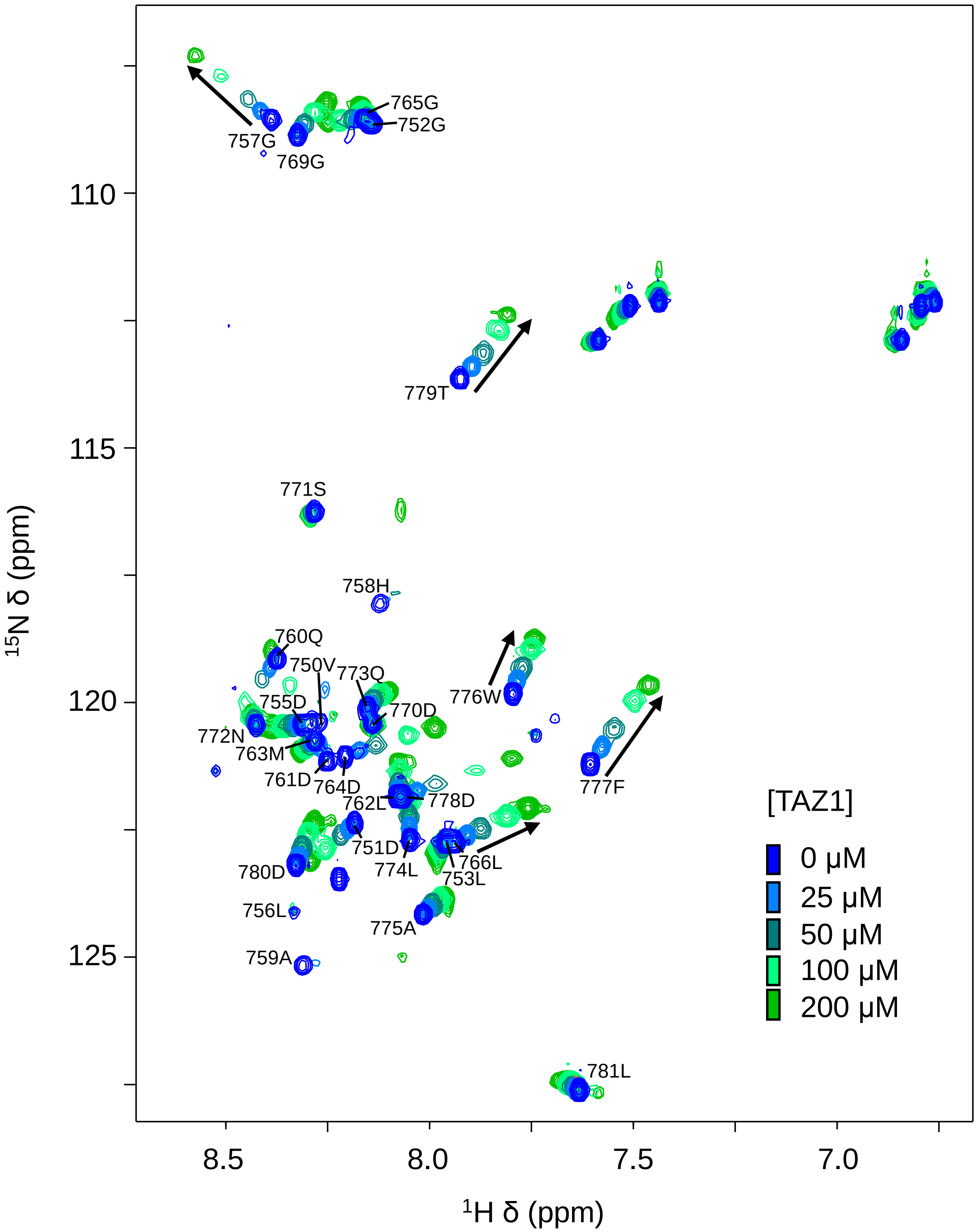


**Fig. S3.** $^{1}$H-$^{15}$N HSQC of 100 μM $^{15}$N-labelled βCAT$_{750\text{-}781}$ (royal blue) titrated with up to 200 μM unlabeled TAZ1 (green). Resonance assignments and direction of peak movement upon addition of TAZ2 are indicated.

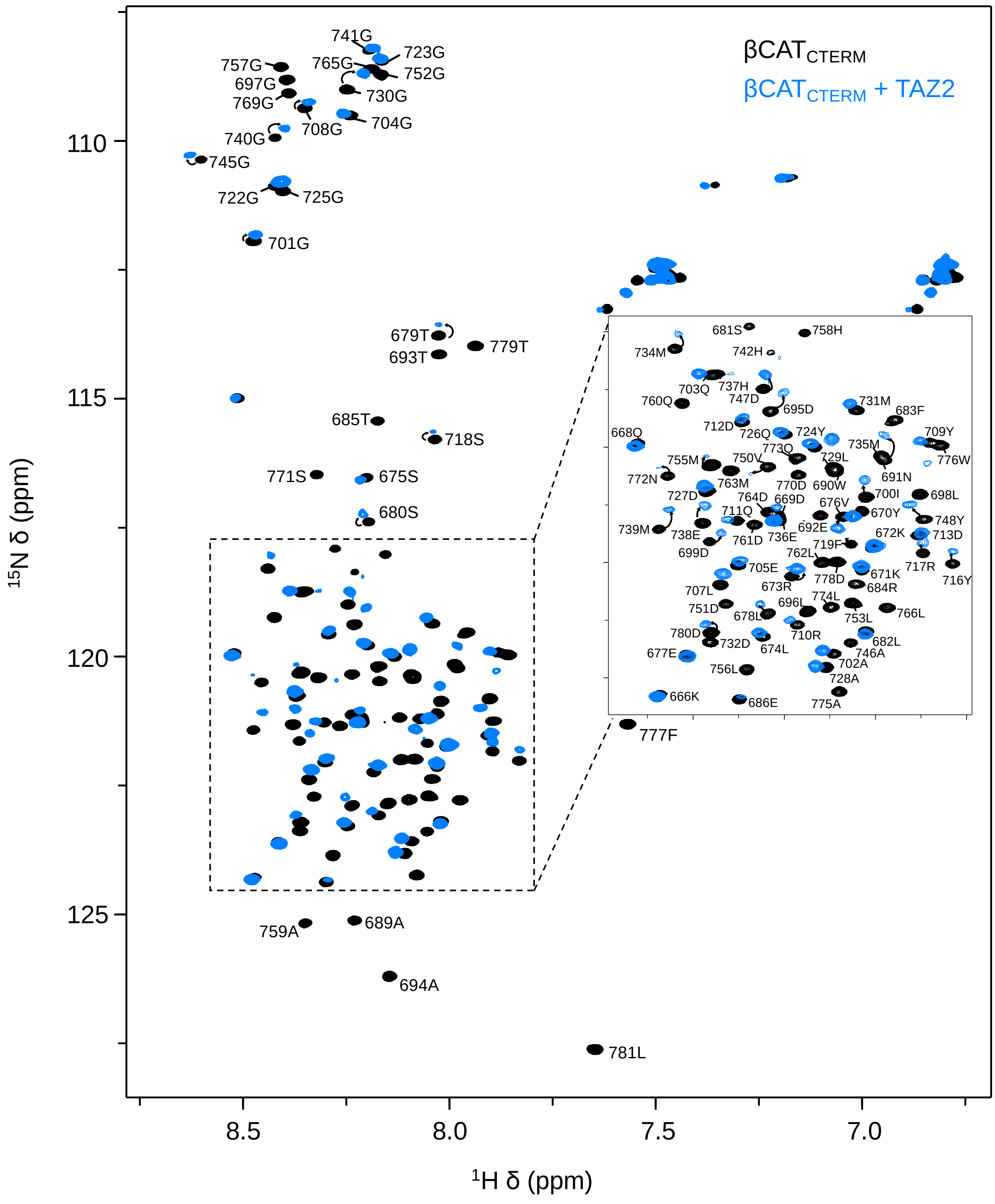


**Fig. S4.** $^{1}H$-$^{15}N$ HSQC of 100 µM $^{15}N$-labelled $\beta CAT_{CTERM}$ overlayed in the absence (black) and presence (blue) of 400 µM unlabeled TAZ2. Resonance assignments and direction of peak movement upon addition of TAZ2 are indicated.

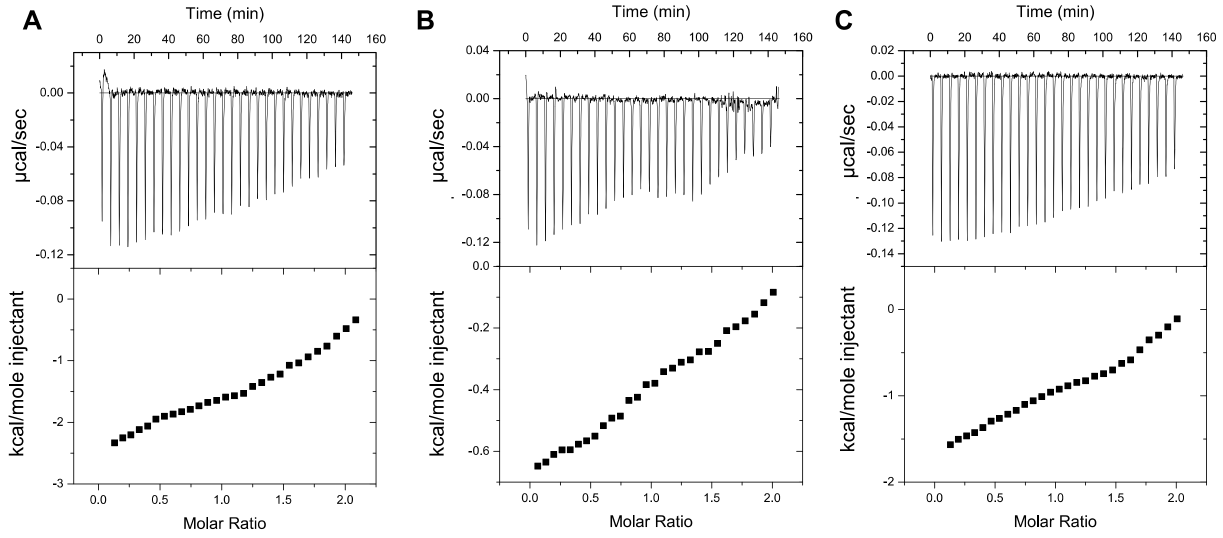


**Fig. S5.** (A) Isothermal titration calorimetry thermograms of 100 µM TAZ1 titrated into 8 µM $\beta CAT_{CTERM\Delta 757\text{-}761}$ (B) $\beta CAT_{CTERM\Delta 761\text{-}765}$ or (C) $\beta CAT_{CTERM\Delta 776\text{-}779}$.
